# 3D Bioelectronics with a Remodellable Matrix for Long-term Tissue Integration and Recording

**DOI:** 10.1101/2022.09.26.509464

**Authors:** Alexander J. Boys, Alejandro Carnicer Lombarte, Amparo Güemes Gonzalez, Douglas C. van Niekerk, Sam Hilton, Damiano G. Barone, Christopher M. Proctor, Róisín M. Owens, George G. Malliaras

## Abstract

Bioelectronics hold the key for understanding and treating disease. However, achieving stable, long-term interfaces between electronics and the body remains a challenge. Implantation of a bioelectronic device typically initiates a foreign body response, which can limit long-term recording and stimulation efficacy. Techniques from regenerative medicine have shown a high propensity for promoting integration of implants with surrounding tissue, but these implants lack the capabilities for the sophisticated recording and actuation afforded by electronics. Combining these two fields can achieve the best of both worlds. Here, we show the construction of a hybrid implant system for creating long-term interfaces with tissue. We create implants by combining a microelectrode array with a bioresorbable and remodellable gel. These implants are shown to produce a minimal foreign body response when placed into musculature, allowing us to record long-term electromyographic signals with high spatial resolution. This device platform drives the possibility for a new generation of implantable electronics for long-term interfacing.

## 1. Introduction

A new generation of bioelectronic technologies provides us with unprecedented means to monitor, stimulate, and understand tissues.^[1]^ A shift to fully flexible electronics allows for maintenance of contact with the body and enhanced recording fidelity.^[2]^ These devices interact with small ionic fluxes that arise from cellular action, e.g. signaling across neuromuscular junctions, to record from or induce a response in cells.^[3,4]^ Recent efforts have been directed at developing technologies for neuroprostheses that control bionic limbs by decoding electromyographic signals.^[5,6]^ Recording devices are typically placed onto the surfaces of tissues to capture cellular signals in their vicinity. Non-invasive electrodes allow for continuous recordings without risk of immune response.^[7]^ However, the majority of information processing performed by tissues occurs within the tissue rather than on its surface, meaning significant information is lost or inaccessible. Use of implantable devices provides a means for reaching this information, but these probes typically cause a foreign body response, associated with fibrotic encapsulation,^[8]^ making these technologies difficult to implement long-term.^[5]^ Small or compliant structures can ameliorate the fibrotic response but do not eliminate it entirely.^[3]^ Efforts to overcome this issue have broadly fallen into two categories, both involving application of coatings to implanted materials for reducing immune response. The first focuses on reducing mechanical mismatch between implantable materials and surrounding tissue.^[9–11]^ This idea has continued to gain precision and complexity through the use of tailored, gel-based coatings.^[12–15]^ The second strategy utilizes pharmacologic means to reduce immune response through sustained release of drugs from coatings on implants.^[16–18]^ However, both methods essentially focus on ‘hiding’ the implant from the body rather than truly integrating the implant with surrounding tissue.

Principles from regenerative medicine provide means to induce integration between implants and the body. These strategies use tissue mimetic architectures, employing gels and/or cells as a modality for repair and regeneration, with recent work focusing on applications in numerous tissues throughout the body.^[19–21]^ These implant systems have been shown to repair damaged tissue,^[22]^ restore native function and mechanics in tissue defects,^[23]^ integrate and interact with local cellular populations, including neurons,^[24]^ or even replace damaged tissues entirely.^[25]^ One unifying aspect for implantable systems in regenerative medicine is the ultimate fusion of the engineered structure with surrounding tissue through the use of 3D tissue-like gels or similar architectures. However, these implants lack further functionality, possessing no externally accessible components. By applying these design strategies to bioelectronics, we can potentially influence the way tissue interacts with bioelectronic devices.

Hybrid fabrication strategies allow us to bring together the functionality of bioelectronics with the biological compatibility and integrative capacity of regenerative medicine. Combination of these approaches provides a means for producing implant architectures that are of similar scale, structure, and composition to local tissue, enabling access to new types of information in the body. Here, we detail the design and implementation of a new implantable platform for integration and recording in tissue. We show that these implants can seamlessly integrate into musculature and record spatially-resolved and high resolution signals to create long-term interfaces between electronics and the body.

## 2. Results

We designed a bioelectronic device for enhanced tissue integration. This device possesses individualized, flexible recording leads designed to mimic tissue structures, such as muscle fibers. We took inspiration from a minimalist design strategy^[26]^ to reduce the amount of material within the insertion site. Our device has 32 independently articulated leads (**Figure 1**a,b), each with approximate 10 × 4 μm cross-sections. These leads separate from the base device (Figure 1c) and terminate in 20 μm diameter recording pads (Figure 1d). The electrodes are arrayed into 8 groups in tetrode-like configurations, where the longest tetrode is 2.25 mm, and the shortest 0.5 mm. The mean impedance at 1000 Hz for electrodes was measured at 48.9 kΩ (Figure 1e, **Figure S1**).

**Figure 1.**
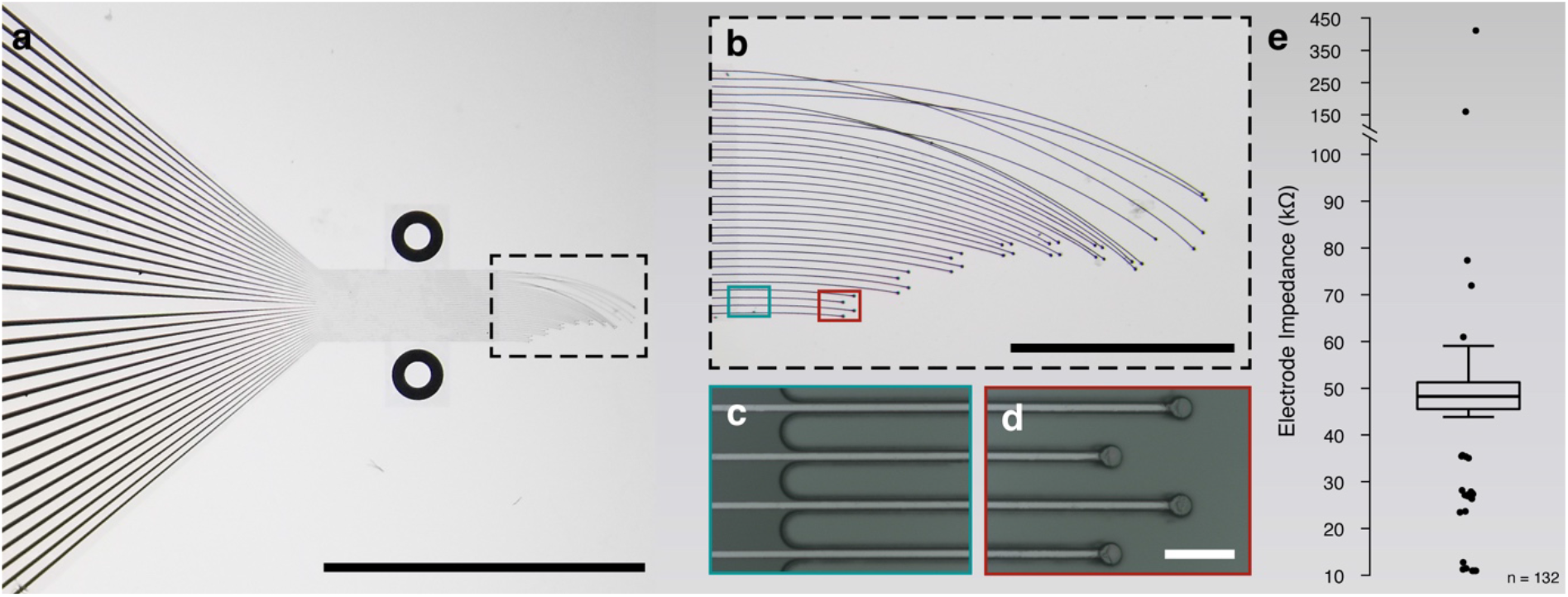
Minimalist bioelectronic device design and implementation for hybrid implant manufacture. **a**, Optical image showing bioelectronic device. Scale bar is 5 mm. Device is constructed from gold insulated parylene-C with PEDOT:PSS-coated gold electrodes. **b**, Image showing individually articulated leads. Scale bar is 1 mm. **c**, Image showing the base of the device where leads break away from the main body of the structure. **d**, Image of the electrodes showing PEDOT:PSS-coated gold electrodes. Scale bar is 50 μm. **e**, Box plot of impedances measured at 1000 Hz using an Intan recording system for N = 5 devices, each containing 32 electrodes, for a total of 160 electrodes. Outliers were defined using the interquartile range and are shown as black circles on the box plot. Broken electrodes (28 of 160), defined as having an impedance >1 MΩ, were excluded from the distribution in this figure, as these electrodes could not be used for recordings, leaving n = 132 electrodes represented in this plot. Supporting information contains a box plot for the full impedance distribution, including broken electrodes.

We utilized principles from regenerative medicine to develop a hybrid implant to promote tissue ingrowth around and through the bioelectronic device architecture (**Figure 2**a). We extracted type I collagen from rat tail tendons for use as a remodellable gel matrix. We injection molded this collagen around the active portion of the implant to create a three-dimensional array of bioelectronic leads within a fibrillar collagen gel (Figure 2b, **Figure S2**), where direct contact between the leads and gel is maintained (Figure 2c). Next, we performed electrical impedance spectroscopy (**Figure S3**) to confirm that the collagen has minimal impact on the recording capabilities of the device. These measurements revealed little difference before and after gel embedding, indicating no inhibition of electrical recording capability (Figure 2d,e, **Figure S4**).

**Figure 2.**
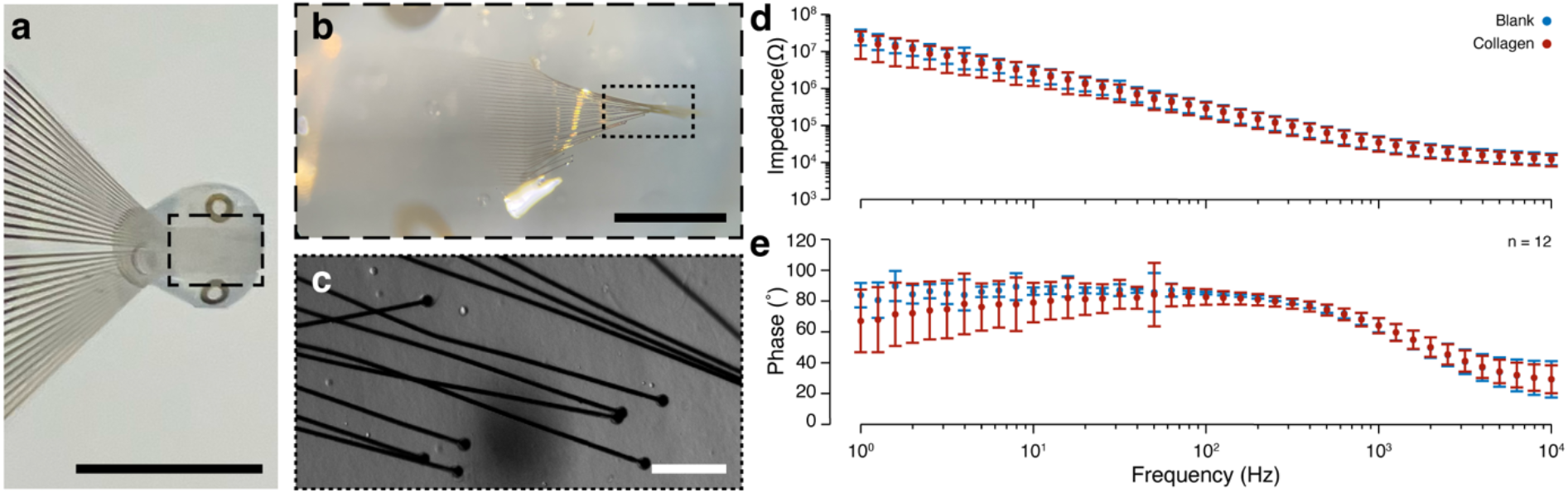
Hybrid implant structure and properties. **a**, Image of hybrid implant with 8 mg/mL fibrillar collagen gel. Scale bar is 5 mm. **b**, Image showing functional end of implant with leads distributed in gel. Scale bar is 1 mm. **c**, Image collected on confocal microscope using Zeiss electronically switchable illumination and detection module (ESID), showing leads embedded in gel. Scale bar is 100 μm. **d**, Bode plot for magnitude and **e** phase of impedance for devices prior to gel embedding and after gel embedding. Data shows the average value +/- the standard deviation for n = 4 arbitrary electrodes on each of N = 3 devices, representing a total of n = 12 electrodes. Broken electrodes were considered to have an impedance of >1 MW and were not measured. For measurement purposes, spectra were recorded for implants without collagen, then the implants were embedded in the measurement dish by dropping pre-gelled collagen below the leads, placing the leads onto the collagen, and then placing more pre-gelled collagen on top of the leads. Collagen was allowed to gel before, and spectra were collected for gel-embedded implants.

First, we sought to understand the degradation and remodeling behavior of collagen in the context of our bioelectronic device architecture using an *in vivo* intramuscular model. We constructed hybrid implants with large collagen gels surrounding the leads, ~1.5 × 1.5 × 5 mm (Figure S2) and placed these implants into dorsal musculature of female Lewis rats. Rats were sacrificed at Days 3, 7, and 14 (n = 2 per timepoint), and tissue was analyzed histologically (**Figure S5**). Implants lacking collagen were placed into a second group of rats with the same timepoints (n = 2 per timepoint) to compare with hybrid implants (**Figure S6**). Cellular infiltration into the periphery of the collagen gel was noted at Day 3, with deeper infiltration evident by Day 7. By Day 14, collagen gels were fully infiltrated with cells, but the gel was still evident. A small number of leads showed evidence of cellular encapsulation by Day 14 for both hybrid and implants lack collagen (Figure S5, Figure S6). The remainder did not show any evidence of cellular encapsulation.

Following successful indications of integration, we performed electrophysiological recordings using a similar intramuscular implantation model. We reduced the implant size to a ~800 μm diameter gel encapsulating the leads and used a wire to assist in implant placement (**Figure S7**). We placed a hybrid implant into a small incision in the dorsal musculature of a rat with a wired connection through an access port on the head (**Figure 3**a-c). We performed recordings at Days 1, 3, and 7 (**Figure S8**) by placing the rat into a clear chamber and allowing the rat to move freely. Video with simultaneous recording shows that movement of the implanted muscle group was associated with increased signal (**Video S1**). Signals were only observed during muscle actuation, regardless of implant conformation (Figure 3a,c), indicating the signal is electromyographic (EMG) in nature. As the animal was able to move freely, the number of events observed during recordings are associated with the rat’s interest in exploring the environment. As such, analysis focused on signal magnitude rather than temporal association. Qualitative analysis of signal shows a decrease in baseline throughout the experiment (Figure 3d-f), along with a monotonic decrease in low-frequency spectral energy across the measured time points (Figure 3g-i). A quantification of the signal-to-noise ratio (SNR) (Figure 3j) showed an increase from Day 1 to 7 but a decrease from Day 1 to 3.

**Figure 3.**
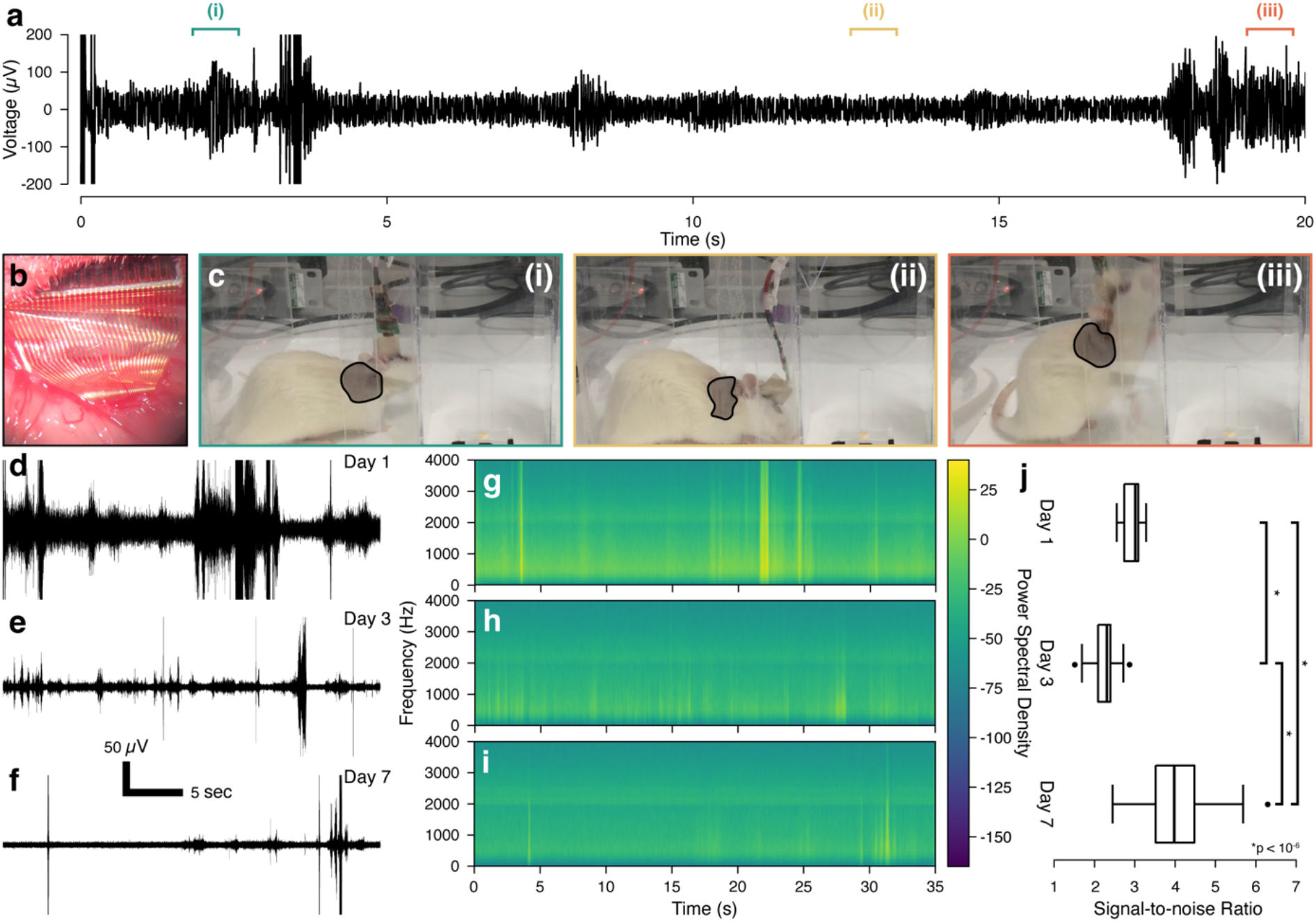
Electrophysiological analysis of hybrid implant recording capabilities in an intramuscular model. **a**, Representative time trace of electrode recordings on Day 1. Trace is bandpass filtered from 400 to 2000 Hz. **b**, Image of hybrid implant inserted into dorsal musculature of a female Lewis rat. **c**, Images showing (i) rat moving her head corresponding to signal in above time trace, (ii) rat sitting still corresponding to baseline in above time trace, and (iii) rat climbing the wall corresponding to signal in above time trace. Highlighted areas show the implant location and approximate conformation of the implanted muscle group. Representative time traces at **d** Day 1, **e** Day 3, and **f** Day 7. Traces are filtered from 400 to 2000 Hz. Spectrograms for **g** Day 1, **h** Day 3, and **i** Day 7. **j**, Box plot for Days 1, 3, and 7 for signal-to-noise ratio (SNR) for all functional electrodes on the implant. The SNR was calculated as the ratio of the mean spike peak-to-peak voltage divided by twice the baseline standard deviation value in a period with no activity.^[27]^ SNR (n = 30 electrodes paired across all days) was found to be significantly different for all time points using a paired t-test with an applied Bonferroni correction for the 3 tests. For Day 1 to Day 3 (p < 10^−9^), for Day 1 to Day 7 (p < 10^−6^), and for Day 3 to Day 7 (p < 10^−14^). Implants were constructed using 8 mg/mL collagen gels.

Next, we performed further analysis on these electrophysiological recordings to understand the type of data that can be collected from implants placed in this manner. An adaptive threshold (**Figure S9**) was applied to identify spikes in the signal for all three days analyzed (**Figure 4**a-c), and the spike rate and mean spike amplitude were calculated for all channels (**Figure S10**). We utilized the tetrode layout of our hybrid implant to examine for spatially-dictated electrophysiological patterns. Comparing time traces of mean spike amplitude within single tetrodes, we found similar waveforms for all days analyzed (Figure 4d-f, **Figure S11**). Comparing mean spike amplitude for electrodes from mixed tetrodes (Figure 4g-i, **Figure S12**), we observed differences in both the amplitude and, in some cases, temporal location of spiking. This trend becomes more evident throughout the time points, with considerable differences in peak location observable by Days 3 and 7 (Figure 4g-i), possibly indicative of individual motor units.

**Figure 4.**
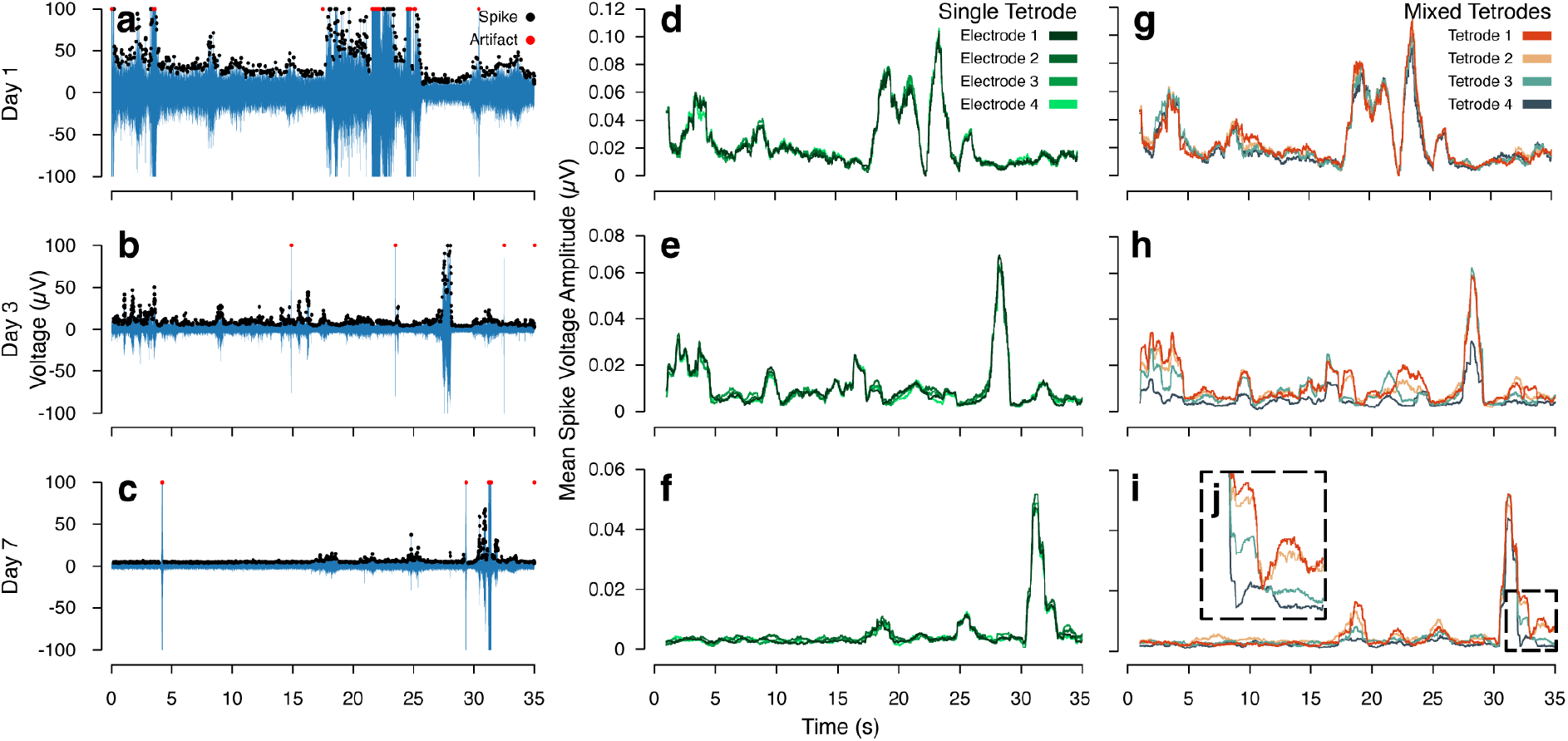
Electrophysiological analysis of recordings from hybrid implant placement. Representative time traces for **a** Day 1, **b** Day 3, and **c** Day 7. Traces are high bandpass filtered from 400 to 2000 Hz. Output voltage traces are shown in blue. Spikes are marked using black circles, and artifacts are marked using red circles. Mean spike voltage amplitude for identified spikes, calculated using a rolling window of 1 s, for four electrodes from a single tetrode at **d** Day 1, **e** Day 3, and **f** Day 7, and for four electrodes, each from a different tetrode, at **g** Day 1, **h** Day 3, and **i** Day 7. **j**, Inset shows differences in spike height and location for an arbitrary portion of this time trace at Day 7. This analysis was conducted to examine for spatial differences in electrophysiological data. As the implants were placed into a singular muscle group, spatial differences at this scale would be indicative of individual motor units. These data are further supported by the evolution of differences in temporal spike location and spike amplitude per trace, as healing and repair occurs from Day 1 to Days 3 and 7.

To understand the likelihood of recording individual motor units, we performed a long-term analysis for the resultant location of implant leads within muscle and their ultimate integration with the body. We constructed implants similar to those used for electrophysiological measurements (Figure S7) and placed these implants into the dorsal musculature of rats (n = 7), as before. We placed an ipsilateral implant (**Figure S13**a) lacking collagen for comparison of encapsulation by fibrotic tissue. Rats were sacrificed after 2 months, by which time chronic foreign body response with associated fibrotic encapsulation would be expected to develop. A gross examination of tissue revealed no large-scale evidence of a fibrotic tissue in the proximity of the implant leads (Figure S13b,c) for all animals. Next, we performed histological analysis to examine for fibrotic encapsulation at the microscale (Figure 5, **Figure S14**). We did not observe collagen gel in any samples (**Figure 5**a), finding only muscle (n = 6). We also located individual leads within the tissue (n = 3 animals), finding them between muscle fibers in all cases for both hybrid implants and ipsilateral controls (Figure 5b,c, Figure S14), corroborating observations from electrophysiological analysis. For hybrid implants, we did not observe any indication of fibrotic encapsulation for two animals, while we did note a minor response for the third animal. For the animal exhibiting encapsulation, the implant leads remain approximately 20 μm from adjacent muscle tissue. For implants lacking collagen, we observed no indication of encapsulation for one animal and more significant fibrotic tissue in two animals, with leads positioned approximately 50 to 100 μm from adjacent tissue (Figure S14). As this study was conducted with ipsilateral controls, the animal with fibrotic encapsulation of the hybrid implant showed this response for the implant lacking collagen (**Table S1**).

**Figure 5.**
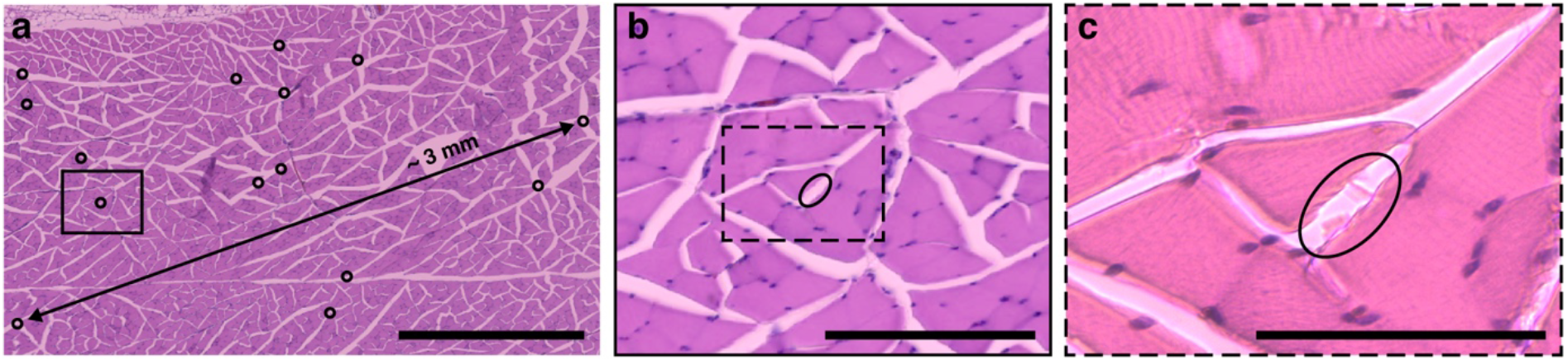
Long-term intramuscular integration of hybrid implants with surrounding tissue. Histological images show the locations of various leads within dorsal musculature after 2 months of implantation (Day 56) in female Lewis rats. **a**, Hematoxylin and eosin staining of implant locations in muscle with inset **b** and inset **c** showing the specific location of two leads. Black circles identify individual lead locations. The longest observed distance between two leads is approximately 3 mm, as indicated. Brightness and contrast were adjusted for **a** and **c** for image clarity. Scale bars are **a** 1 mm, **b** 200 μm, and **c** 100 μm. Collagen gels were made at 10 mg/mL.

## 3. Discussion

We demonstrate a new system for long-term integration of bioelectronic sensors with tissue. The combination of bioelectronics with principles from regenerative medicine creates a customizable and functional platform for generating stable interfaces with the body. Our hybrid implants use a minimalist device structure with high efficacy long-term recording. This design in combination with the use of remodellable collagen gels promotes the spread and integration of these devices into tissue capabilities to maximize cellular access for the implant while minimizing the foreign body response. We were able to use these devices to record high resolution EMG signals with increasing SNR over time, which appear to be derived from individual motor units.

Hybrid fabrication strategies allowed us to produce architectures similar in size and structure to tissues, while also providing three-dimensionality to microfabricated devices. We realized this methodology with small, individually articulated leads (Figure 1) and a remodellable gel (Figure 2). This implant design provides multiple advantages over traditional implantables. First, while flexible electronics are typically used to reduce the body’s response to implanted materials,^[2,3,28,29]^ the compliance of these devices makes placement and maintenance of their position in the body challenging. Our use of injection molding gives us control over the resultant geometry of the implant (Figure S2, Figure S7), allowing us to set the location of the electrodes in 3D space during implantation. Secondly, while application of gel coatings to bioelectronics is a well-known means to improve mechanical interfacing,^[12–15]^ coatings primarily hide the implant from surrounding tissue rather than inducing integrative scenarios. Our design creates increased opportunity for tissue ingrowth through the implant architecture using a 3D bioresorbable gel while still providing options for mechanical tuning of the implant properties for interfacing purposes. Our implant construction method also allows for inclusion of cells in this 3D matrix. We demonstrated this capability *in vitro* (**Figure S15**), forming gels with both stem cells and neural cell types. We chose not to include cells in our study, given the rapidity with which we observed cellular infiltration into our hybrid implants (Figure S5). Various researchers have proposed the development of cellularized bioelectronics for *in vivo* usage,^[30,31]^ but the studies which have implanted these types of structures used stiff devices with associated cells,^[32,33]^ which did not eliminate the body’s immune response. These studies also focused on the brain and did not record electrophysiological signals. Other studies have constructed nerve conduits^[34–36]^ or tissue engineered neural structures^[24]^ but did not focus on tissue integration or electrical functionality. Regardless, while we did not require cellularization to drive tissue integration with our hybrid implants, we could envision inclusion of cells for producing larger implantable structures or modified cells for immunotherapies or similar.

Our implant construction was designed to reduce the foreign body response by improving integration and repair. After 2 months of implantation in muscle, we observed either no or minimal fibrotic encapsulation for our hybrid implants (Figure 5, Figure S14). Even in cases of observed encapsulation, the implanted leads remain approximately 20 μm from surrounding tissue at 2 months, meaning recording through this small fibrous layer is very plausible. In implants lacking collagen, we observed more severe fibrotic encapsulation, evidenced by the larger size of surrounding fibrotic tissue (Figure S14). As collagen can be remodelled and degraded, the hybrid implants’ construction may have sufficiently slowed the body’s reaction to the bioelectronic device. While this result is very promising, more analysis with larger numbers of animals needs to be conducted to determine specific drivers for resultant foreign body response. We considered quantification of un-encapsulated versus encapsulated leads, finding for example 22 to 11 leads, respectively, for hybrid implants. However, given the tortuous nature of the implanted leads, we cannot confirm if we are double counting leads in any case. As such, we chose not to pursue this quantification further. Regardless, we speculate that the body possesses an upper size limit for the foreign body response between 1 and 10 μm. Similar size limits have previously been proposed,^[37]^ but testing these limits is challenging. One study examined foreign body response as a function of implant size, finding that it was continuously reduced down to 6 μm but was still apparent.^[38]^ However, as size is only one driving factor,^[29]^ direct comparisons are likely impossible. Regardless, a greater understanding of constraints that can be applied to eliminate the foreign body response would be extremely informative for future implant designs.

We found that our hybrid implants were ultimately located in the interstitial space between muscle fibers (Figure 5c). We speculate that this placement may be driven by mechanical actuation of muscle and the resultant orientation caused by continuous application of mechanical force.^[39]^ This result provides us with some interesting opportunities to electrically access portions of muscle that are unreachable with other implant designs. Most intramuscular EMG recordings are performed with electrodes that are at least 1 to 2 orders of magnitude larger than our electrodes.^[40–44]^ Studies focused on high resolution EMG, with electrodes of the same order of magnitude as ours, almost universally utilize needles,^[45]^ limiting their long-term feasibility. The few studies that have approached intramuscular^[46]^ or similar^[47]^ recordings with array-based microelectrodes have observed electrical behavior associated with different neuromuscular units, finding changes in spike timing, etc. No implants, of which we are aware, have been able to successfully integrate with tissue to the degree necessary for long-term intramuscular recordings, providing little comparison for interpreting our data. We show an overall increase in SNR between Days 1 and 7 of implantation. We observed a decrease in SNR at Day 3, which may have been related to high degree of activity by the animal on this day, as the baseline appears lower despite the apparent drop in SNR. However, the SNR was shown to be significantly above Days 1 and 3 on Day 7, indicating improved integration between the implants and surrounding tissue, as is corroborated by our long-term histological analysis.

We used the tetrode-like construction of our implants to understand the origin of our recorded data. Throughout the experiment, electrodes within defined tetrodes show similar spiking (Figure 4d-f). However, when comparing electrodes from mixed tetrodes, we note temporal differences in peak location and mean spike amplitude (Figure 4g-i), which are not evident until Days 3 and 7. One possibility is that these changes are associated with EMG from adjacent and separate muscle groups. As we do not observe these differences on Day 1, this is unlikely, since these signals would be traversing tissue in a similar manner to non-invasive recordings, which would be feasible regardless of the integrative state of local tissue around the implant. A second possibility is that these changes are due to innervation of a single muscle group by multiple motor units. This possibility accounts for both changes in peak amplitude and peak location. First, a given electrode would be located at a different relative distance to a motor unit versus any other electrode, accounting for changes in peak amplitude. Second, as the muscle heals from implantation surgery, other motor units located farther from the site of implantation could become electrically accessible, accounting for changes in peak location. A limitation to this device design is that, as the electrodes are individually articulated, we cannot assume that any electrode remains in its exact tetrode conformation. However, as electrodes in different tetrodes are of the same approximate length and as we observe conservation of signal waveform within tetrodes, we believe we are most likely recording different motor units with our implant configuration.

We envision these devices as useful for applications in neuroprostheses. For example, targeted muscle reinnervation^[48,49]^ has been shown to be an effective means for providing neurally-driven actuation of prostheses. In this procedure, nerve endings from amputated arms are sutured into the pectoral muscles of the patient, allowing the nerves to reinnervate these muscles. Subsequent neural impulses through these nerve endings result in EMG signals, which can be recorded using cutaneous electrodes.^[48,49]^ As we show above, our hybrid implants are able to infiltrate into musculature, providing a means for high resolution recording. These implants could be placed with the nerve stump into the muscle to monitor and record muscle reinnervation. The spread of implant leads within musculature would provide a means to record from the different aspects of the nerve stump as it grows into and reinnervates the muscle. This type of therapeutic could offer a method for driving a neuroprosthetic with high efficiency and high resolution inputs.

While we were able to show successful integration and recording in muscle, we foresee these implants as useful in other regions of the body as well. Collagen or gelatin has previously been shown to improve integration and recovery in the brain as a coating on non-functional probes^[50]^ or as a free-standing matrix without electronics.^[51]^ As a preliminary look at applications in the central nervous system, we placed these hybrid implants into the brain (Supporting Methods, **Figure S16**, **Figure S17**, **Figure S18**, **Figure S19**). While our gel-based modality is different from previous designs in literature, we also noted a limited immune response for our hybrid implants. This immune response was similar to contralaterally-placed implants lacking collagen (Figure S16, Figure S17). This finding is potentially substantial in that the implants lacking collagen are much smaller (50 μm versus ~800 μm diameter). We also observed cellular infiltration into hybrid implant interior (Figure S16), which is promising, as this is indicative of a repair mechanism. Regardless, refinement in fabrication to reduce the size of our implants would make these devices suitable for many applications in the brain. However, areas like traumatic brain injury, which produce fissures in the brain, are promising even for the current design. Previous studies have shown that scaffold or gel-based approaches can help treat traumatic brain injuries,^[52,53]^ while other studies utilized electrical stimulation to reduce neural degeneration at the injury site.^[54,55]^ Our hybrid implants could simultaneously provide both options to assist in mitigation and repair of these injuries. Similar strategies could also be applied to other tissues in the body, e.g. to increase recovery rates for patients with severe orthopedic injuries by monitoring and stimulating tissue to promote regrowth.

As we look forward, we must consider translation of these implants to humans. For our hybrid implant construction, we utilized Lewis rat collagen as an implant material in Lewis rats. This strategy allowed us to focus on integration and foreign body response rather than immune response and rejection. To move towards human usage, we would likely need to change the matrix material to a more suitable candidate. Various companies are focused on the development of biomaterials for regenerative medicine. Notably, more companies in this field are focused on biomaterials development than any other area.^[56]^ As tissue ingrowth is a cornerstone of regenerative medicine, the hybrid fabrication strategies we show here are applicable to many of these developing technologies, providing a pathway for translation.

## 4. Conclusion

Overall, we show that hybrid fabrication approaches can be used to produce functional bioelectronic devices that promote tissue ingrowth and long-term integration. We produced a hybrid implant by combining microfabrication with principles from regenerative medicine, while maintaining the overall functionality of the device. We show that these devices can be placed into multiple tissue systems and used to induce long-term integration in muscle without initiating encapsulation by fibrotic tissue. We also showed that these implants can record intramuscular EMGs with high spatial resolution and that the efficacy of these recordings improves over time. These devices provide a new set of tools and a new platform from which to build long-term electronic interfaces between the body and bioelectronics.

## 5. Methods

### Materials

PEDOT:PSS (Clevios PH 1000) was obtained from Heraeus (Hanau, Germany). Ethylene glycol, dodecyl benzene sulfonic acid (DBSA), 3-glycidyloxypropyl trimethoxysilane (GOPS), and poly(ethylene glycol) (PEG) (BioUltra 8,000) were purchased from Sigma Aldrich (St. Louis, MI, USA). Filters for PEDOT:PSS filtration were purchased from Sartorius (Göttingen, Germany). AZ nLOF 2035 photoresist, AZ 10XT 520cP photoresist, AZ 726 MIF developer, AZ 400K developer, and TechniStrip^®^ NI555 were obtained from Microchemicals GmbH (Ulm, Germany). Micro-90 Concentrated Cleaning Solution was obtained from Cole-Parmer (Vernon Hills, IL, USA). Dichloro-p-cyclophane, used for parylene-C chemical vapor deposition (CVD) was purchased from Specialty Coating Systems (Indianapolis, IN, USA). Anisotropic conductive film (ACF) (5 μm particulate) was purchased from 3TFrontiers (Singapore). Flat flex cables were purchased from Mouser Electronics (Mansfield, TX, USA). Microwires (50 μm diameter tungsten wire) for surgical placement and implant fabrication were obtained from the California Fine Wire Company (Grover Beach, CA, USA). Tygon^®^ tubing (0.79mm ID) was purchased from Tygon (Akron, OH, USA). Acetic acid was purchased from Acros Organics (Geel, Belgium). Isoflo® isofluorane and Rimadyl (50mg/mL stock), used for surgical anesthesia and analgesia, were purchased from Zoetis (Parsippany-Troy Hills, NJ, USA). Metacam® Meloxicam (1.5mg/mL stock) for analgesia was purchased from Boehringer Ingelheim Ltd, Animal Health (Ingelheim, Germany). Anti-GFAP (ab7260), anti-CD11b+CD11c [OX42] (ab1211), donkey anti-rabbit IgG H&L (Alexa Fluor® 555) (ab150074), donkey anti-mouse IgG H&L (Alexa Fluor® 488) (ab150105), and H&E staining kit (ab245880) were obtained from Abcam (Cambridge, UK). DAPI staining solution (1mg/mL), LIVE/DEAD^™^ viability/cytotoxicity kit, and EDTA were obtained from Thermo Fisher (Waltham, MA, USA). Dulbecco’s modified Eagle medium (DMEM) (high glucose), nutrient mixture F-12 Ham (F-12), minimum essential medium (MEM) Eagle, fetal bovine serum (FBS), and MEM non-essential amino acid solution (100x) were purchased from Merck (Darmstadt, Germany). L-glutamine (200 mM), penicillin (10,000 Units/mL) streptomycin (10,000 μg/mL) solution, 0.25% trypsin-EDTA, and GlutaMAX^™^ (100x) were purchased from Thermo Fisher (Waltham, MA, USA).

### Microfabrication

Bioelectronic devices were fabricated using photolithographic procedures for production of flexible electronics.^[57]^ Si wafers were used a base substrate for fabrication. Parylene-C was deposited, via CVD, at a 2 μm thickness onto Si wafers using a Specialty Coating Systems (SCS) Labcoater (Specialty Coating Systems – Indianapolis, IN, USA). AZ nLOF 2035 photoresist was used in conjunction with AZ 726 MIF developer for gold patterning. A 10nm Ti adhesion layer, followed by a 100nm Au layer was deposited using a Lesker e-beam Evaporator (Kurt J. Lesker Company – Jefferson Hills, PA, USA). Gold liftoff was performed using Technistrip® NI555. A second 2 μm layer of parylene-C was deposited on top of the gold layer, producing insulated gold tracks. Micro-90 detergent was spincoated onto the wafers to act as an anti-adhesive layer for production of PEDOT:PSS coated electrodes. A further 2 μm of parylene-C were deposited on top of the anti-adhesive layer. The electrodes and connector array for the device were patterned using AZ 10XT 520cP photoresist with AZ 400K developer. Wafers were placed into a PlasmaPro 80 Reactive Ion Etcher (RIE) (Oxford Instrument – Abingdon, UK) to etch out the electrodes and connector array pads. The devices were etched using 8sccm CF_4_, 2sccm SF_6_, and 50sccm O_2_ at 60mTorr. The electrodes were coated with a PEDOT:PSS coating, consistent of 5% (v/v) ethylene glycol, ~30 μL of DBSA, 1% (v/v) GOPS, remainder PEDOT:PSS. This solution was filtered prior to application to the devices. Coatings were spincoated in 3 applications with a heating cycle after each application. The anti-adhesive parylene-C layer was peeled off, leaving PEDOT:PSS only on the electrodes. The devices were heated further to cross-link the PEDOT:PSS coatings. Next, AZ 10XT 520cP photoresist and AZ 726 MIF developer were used to pattern the outlines of the devices. A MA/BA6 mask aligner (Süss Microtec – Garching, Germany) was used for all photolithography. The device outlined were etched using a RIE, and the devices were soaked in water overnight before removal from the Si substrate. A flat cable was bonded onto each implant using a Finetech Bonder Fineplacer® System (Finetech GmbH – Berlin, Germany) to connect the electrodes for recording. In some cases, dummy devices, which lack gold and PEDOT:PSS were used for biocompatibility studies. These devices were prepared in the same manner but only by etching the outline into 4 μm of parylene-C on a Si wafer.

### Collagen Extraction

Collagen was extracted from Lewis rat tails, as previously reported.^[58]^ Briefly, rat tails were excised from rat cadavers and skinned revealing tendons running the length of the tail. Vertebrae were dislocated and removed, pulling each tendon away from the tail. Tendons were removed from vertebrae and placed into 70% ethanol. Tendons are moved into 0.1% (v/v) acetic acid at 150 mL/g of tendon and left to solubilize for 48 hours. This mixture was centrifuged at 9000 RPM, and the supernatant was removed and lyophilized, producing dry collagen. Collagen was reconstituted at 20 mg/mL in 0.1% (v/v) acetic acid for later use.

### Implant Fabrication

Implants were fabricated by placing bioelectronic devices onto a plate or into Tygon® tubing for molding. For implantation, devices were first adhered to a 50 μm diameter tungsten wire for ease of placement using a 12% (w/v) solution of poly(ethylene glycol) in water. To apply collagen gel, collagen was neutralized using a working solution consistent of phosphate buffered saline and sodium hydroxide causing the collagen to form a fibrillar gel.^[58,59]^ This gelling solution was injected around the implants using a syringe. Resultant gels were made at either 8 mg/mL or 10 mg/mL final concentration of collagen.

### Electrical Measurements

The (small signal) impedance spectrum of each probe on the device was measured using an Autolab PGSTAT204 potentiostat (Metrohm – Herasau, Switzerland). The measurement consisted of a platinum mesh counter electrode (Goodfellow – Huntingdon, UK), an Ag/AgCl reference electrode (redoxme AB – Norrköping, Sweden), and the test device submerged in PBS. The test devices were connected to the potentiostat using a custom printed circuit board to route individual electrodes to a pin. The reference electrode was positioned in between the counter electrode and the device. Before performing the EIS sweep, the open circuit potential (OCP) was determined via a search optimization algorithm on the instrument. The search duration was 50 s, with a 0.1 s interval, and the final value was averaged over 10 s. To prevent de-doping of the PEDOT coating on the electrodes, the OCP was constrained to positive potentials. Once the OCP was determined, the EIS sweep was conducted with the OCP as the DC operating point. To perform the measurement, the above 3 electrode setup was used. Impedances were scanned between 1 Hz and 10 kHz with 10 samples per decade at a sine amplitude of 0.01 V_rms_ with a maximum integration time of 0.125 s. These parameters were applied using Nova 2.1.4 software (Metrohm – Herasau, Switzerland). To compare electrode EIS to electrode EIS embedded in collagen, collagen gels were injected directly into the measurement dish around the electrodes at 8 mg/mL collagen gel density. The gels were allowed to form for approximately 30 minutes, and the measurements were repeated as above.

### Animal Surgery

All experimental procedures were performed in accordance with the UK Animals (Scientific Procedures) Act, 1986 and were approved by the animal welfare and ethical review body at the University of Cambridge. Female Lewis rats (200 – 250g), purchased from Charles River Laboratories (Kent, UK), were used for all animal work. Animals were anesthetized for surgery using an isoflurane inhalant. Rats were given a pre-operative analgesic (Rimadyl) along with a saline injection for fluid replacement. After surgery, rats received a post-operative analgesic (Meloxicam) at days 1 and 2, post-surgery. Implants were placed into the dorsal musculature. The implant leads were inserted into the musculature, and the back end of the implant was sutured on top of the muscle. For Day 3, 7, and 14 timepoints, a singular rat received either a hybrid implant or an implant lacking collagen for a total of n=2 rats per timepoint per group. For 2-month time points, rats received hybrid implants with a second implant lacking collagen placed ipsilaterally in a total of 7 rats. Dummy implants, lacking gold with PEDOT:PSS coatings, were used for these time points.

### Electrophysiology

For electrophysiological EMG recordings, an implant was placed into the dorsal musculature as above. In this case, wires connected to the implant were run subcutaneously to an access port on the animals head. The access port was a custom 3D print to facilitate recordings. Recordings were conducted by connecting the implant through a custom printed circuit board to an Intan 32-Channel Stim/Recording Headstage (Intan Technologies – Los Angeles, CA, USA). Data was collected using an Intan RHS Stim/Recording System (Intan Technologies – Los Angeles, CA, USA). Recordings were performed at Days 1, 3, and 7 at a sampling frequency of 30 kHz. Electrophysiological recordings were performed inside a transparent acrylic box (300 × 300 mm), otherwise animal was group-housed with *ad libitum* access to food and water between sessions.

### Electrophysiological Data Processing

Raw signals were first filtered with a Butterworth 4^th^ order bandpass filter with cut-off frequencies at 200 Hz and 2 kHz. To identify high amplitude high-frequency artifacts from the filtered signal, an Envelope Derivative Operator (EDO) signal was implemented to estimate frequency-weighted instantaneous energy of the signal.^[60]^ Peaks over 3 times the standard deviation from the mean of the EDO signal were identified as artifacts. Events of interest (‘spikes’) where identified using two methodologies that were compared. In the first methodology, a constant threshold was calculated and applied to each of the channels *(i)*. The threshold was calculated as 3 times the standard deviation.^[61]^ The amplitude of the neural recordings is typically not stable over time due to changes in the impedance of the electrode-tissue interface. For this reason, an adaptive thresholding was implemented based on the smallest of constant false-alarm rate (SO CFAR) filter,^[62]^ which was has been used in the past to accommodate for the respiratory modulation on vagus nerve recordings.^[63]^ CFAR filters use a sliding window to continuously estimate the background mean power of a signal so that a threshold that maintains a constant false-alarm rate on average can be applied on a per-window basis. This method allows the threshold to adapt to changes in the background mean power. The parameters of the threshold were the window duration (5000 samples on each side), the guard cell duration (50 samples on each side), and the threshold level (3 SDs from the mean). These parameters were heuristically chosen based on empirical results to guarantee a tradeoff between threshold adaptation to sharp changes and peak detection. Peaks above the threshold in each case were extracted, and those peaks within a 0.2 s window centered around an artifact peak were excluded. Finally, a sliding window of 1 second was used to compute the spike rate and the mean amplitude of the peaks within each window. Plots were created to visualize the evolution of these metrics across channels grouped based on their tetrode distribution within the device. The signal-to-noise ratio (SNR) was calculated as the ratio of the mean spike peak-to-peak voltage divided by twice the baseline standard deviation value in a period with no activity.^[27]^

### Histology

For histological analysis, samples were fixed in 4% paraformaldehyde for 48 hours. For paraffin sectioning, samples were transferred to 70% ethanol, paraffin, embedded, and sectioned at 4 μm thickness. Resultant slides were stained using hematoxylin and eosin. For cryo-sectioning, samples were transferred into 30% sucrose in phosphate buffered saline for 72 hours. Some samples required decalcification due to embedded bones in the tissue block. These samples were decalcified as previously described,^[64]^ by applying a 9.5 (w/v%) EDTA in PBS solution (pH = 7.4) for up to 5 days. Samples were sectioned at 12 μm thickness. Samples were either stained using H&E or prepared for immunofluorescence staining. For immunostaining, samples were blocked with 5% donkey serum in phosphate buffered saline for 1hr. Slides were incubated with primary antibodies in 5% FBS in PBS overnight at 4°C. Slides were washed and incubated with secondary antibodies for 3 hours. Slides were washed and stained with DAPI. For 2-month timepoints, tissue from one rat was dissected under a microscope to assist in locating implants and to ensure successful processing for histology. Tissue from the remaining six rats was processed for histology.

### Imaging

White light images were collected using a Thermo Fisher Evos XL (Thermo Fisher – Waltham, MA, USA), a Zeiss AX10 Lab A1 Microscope (Zeiss – Oberkochen, Germany), an iPad Mini 2, and a iPhone 12 mini (Apple – Cupertino, CA, USA). Confocal microscopy was performed using a Zeiss LSM 800 Confocal Microscope (Zeiss – Oberkochen, Germany). Histological images were collected using a Zeiss Axioscan 7 (Zeiss – Oberkochen, Germany).

### Design & Analysis Software

Initial designs for photomasks were constructed in CleWin 5 (WieWeb Software – Hengelo, Netherlands). Photomasks were purchcased from MicroLithography Services (Chelmsford, UK). Confocal and histological images were analyzed in Zeiss Zen 3.1 (Zeiss – Oberkochen, Germany). Graphing and statistics were performed using MATLAB 2019a (Natick, MA, USA). Figures were constructed using Adobe Illustrator 2022 (San Jose, CA, USA).

## Acknowledgments

The authors would like to acknowledge Nicholas Melosh for his advice relating to some of the initial design work for these implants. The authors would like to acknowledge Santiago Velasco Bosom and Ruben Ruiz Mateos Serrano for assistance in developing a method for measuring the electrical impedance spectra of devices. The authors would also like to acknowledge Chiara Barberio and Yash Mishra for providing SH-5YSY cells for implant fabrication. The authors would like to acknowledge the University of Cambridge Institute of Metabolic Science Histopathology Core and the National Health Service (NHS) Human Research Tissue Bank for paraffin processing of samples. This work was partially supported by the ECH2020 FUTURE & EMERGING TECHNOLOGIES (FET) projects BrainCom (732032) and MITICS (964677). This work was also supported by the United States (US) Air Force Office of Scientific Research (AFOSR) under award number FA8655-20-1-7021. A.J.B. would like to acknowledge support from his Cross-disciplinary Fellowship (Grant No. LT000034/2020-C) from the Human Frontier Science Program (HFSP) Organization. A.C.L. acknowledges support from the University of Cambridge for a Borysiewicz Interdisciplinary Fellowship and the Wellcome Trust for a Junior Interdisciplinary Fellowship. A.G.G. acknowledges funding support from the Royal Commission for the Exhibition of 1851. D.C.v.N. would like to acknowledge support from the W.D. Armstrong Trust Fund and the Oppenheimer Memorial Trust. D.G.B. would like to acknowledge support from the UK Health Education and National Institute for Health Research (HEE/NIHR Integrated Clinical Academic Program Clinical Lectureship CL-2019-14-004).

