## Supporting Information for "3D Bioelectronics with a Remodellable Matrix for Long-term Tissue Integration and Recording"

<u>Supporting Methods</u>

*Brain Surgery* All animal work was licensed and performed as permitted under UK law. The association cortex was chosen as an implantation site to minimize potential complications in analysis of immune response. Other areas of the cortex, e.g. the sensory motor cortex, can have significant negative impacts on rodent behavior, such as autophagia. For surgeries, procedures were performed using a stereotactic frame. Two craniotomies were performed to place two implants contralaterally. Hybrid implants, ~800 mm in diameter (Figure S7), were placed into one side of the brain using a 50 mm tungsten wire as a shuttle. Implants lacking collagen gels were placed contralaterally as a comparison for immune response to collagen gels. Animals were anesthetized for surgery using an isoflurane inhalant. Rats were given a pre-operative analgesic (Rimadyl) along with a saline injection for fluid replacement. After surgery, rats received a post-operative analgesic (Meloxicam) at days 1 and 2, post-surgery. Rats were sacrificed after 3 days, and the implant sites were examined using immunofluorescence for activated microglia and astrocytes (Figure S17, Figure S18, Figure S19, Figure S20). We observed cellular infiltration into the hybrid implant interior by activated microglia and astrocytes at Day 3 (Figure S17a-c). Activated microglia were not evident outside the immediate area of the implant (Figure S17d), indicating that the implants do not produce a long-range immune reaction. Implants lacking collagen showed a similar immune response (Figure S18), with activated microglia becoming less evident at a similar distance from the implant as was observed for hybrid implants.

*Cell Extraction* Bone marrow stromal cells were extracted as previously described.[58,64–66] Briefly, femurs and tibias were dissected from rats. The epiphyses of these bones were removed, and bone marrow was flushed out of the bones using a syringe. This suspension was passed through a cell strainer (70 mm) and spun down at 200xg at 4˚C for 5min. These cells were re-suspended in fresh media (DMEM with 10% FBS, 1% P/S, and 2mM L-glutamine) and plated into Petri dishes for culture. Media was changed over the first 3 days to remove non-adherent cells. Resultant cell populations were frozen for later use.

*Cell Culture* Bone marrow stromal cells were cultured in growth media (DMEM with 10% FBS, 1% P/S, and 2mM L-glutamine) with media changes every 2-3 days. SH-SY5Y cells were cultured in neural growth media (1:1 MEM:F12 with 10%FBS, 1% non-essential amino acids, 1% GlutaMAX, and 1% P/S) with media changes every 2-3 days. Cells were passaged using trypsin.

Supporting Figures

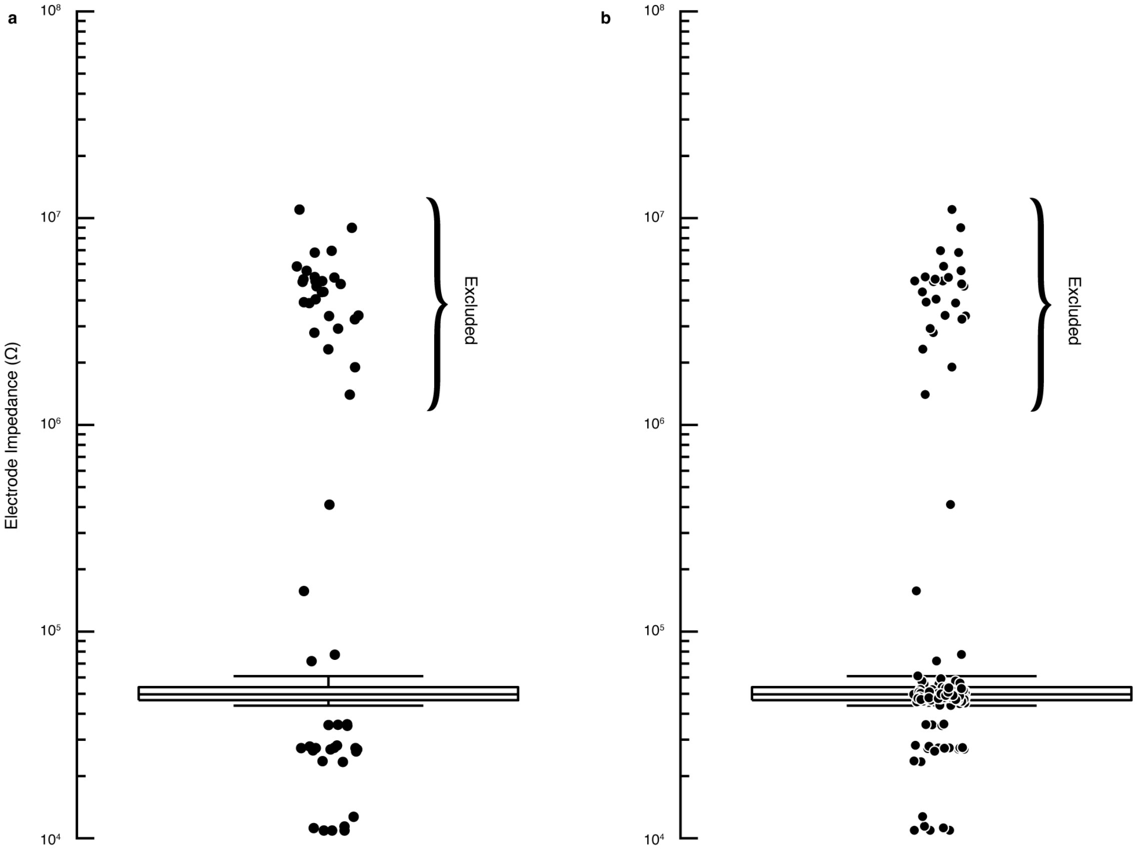


**Figure S1.** Distribution of impedances for all electrodes. **a**, Box plot of impedances for all electrodes. Outliers are shown as black circles. Bracket indicates broken electrodes, which were excluded from box plot in Figure 1. Outliers were defined using the interquartile range. **b**, Box plot of impedances for all electrodes with all points plotted as black circles over box plot.

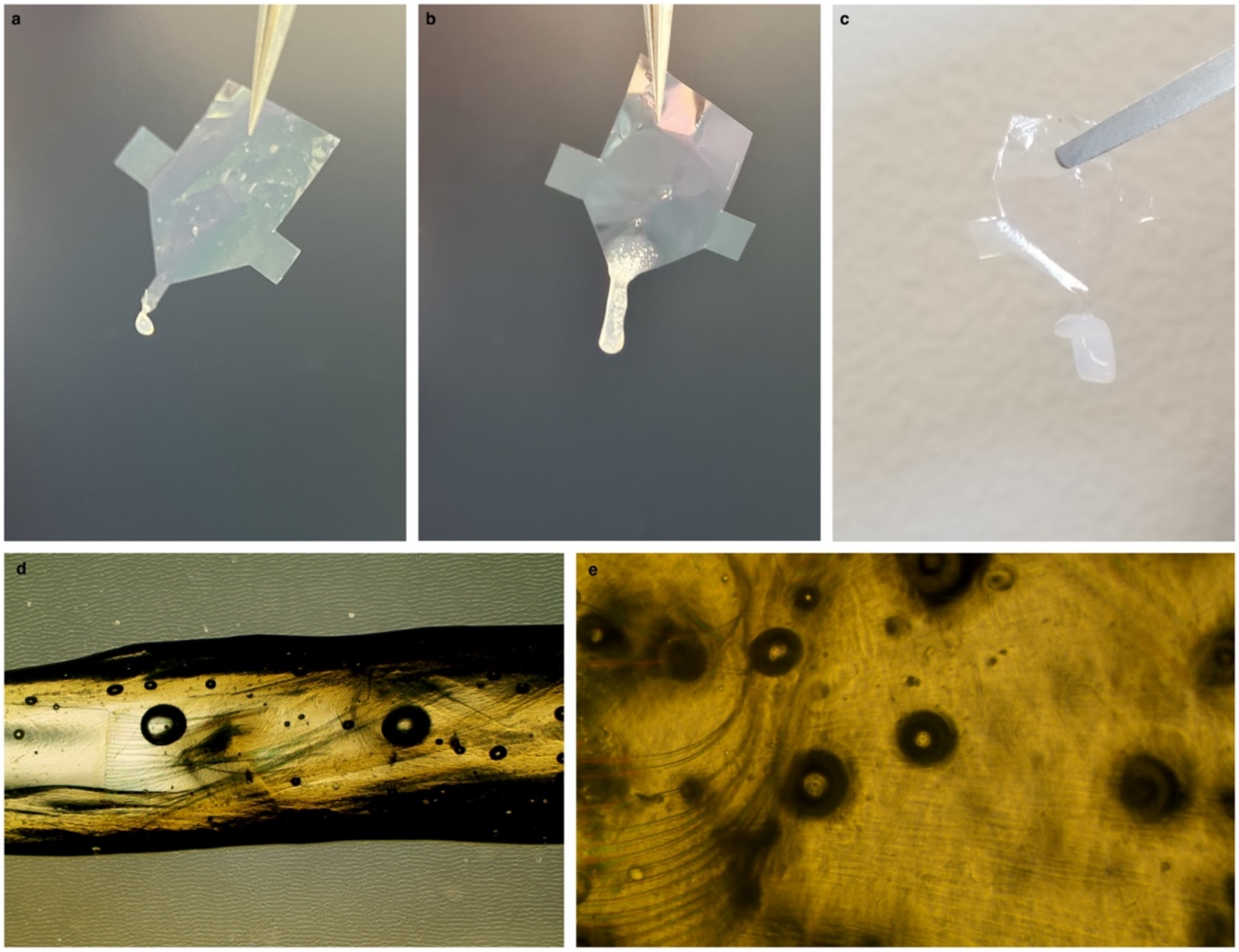


**Figure S2.** Image of dummy implants (lacking gold and PEDOT:PSS components, showing structure of hybrid implants. **a**, Image showing implant with small gel. **b**, Image showing implant wither intermediate-sized gel. **c**, Image showing implant with large gel. Images **d** and **e** show examples of leads from dummy implants distributed in 3D inside of collagen gels. The black circles are bubbles in the gel.

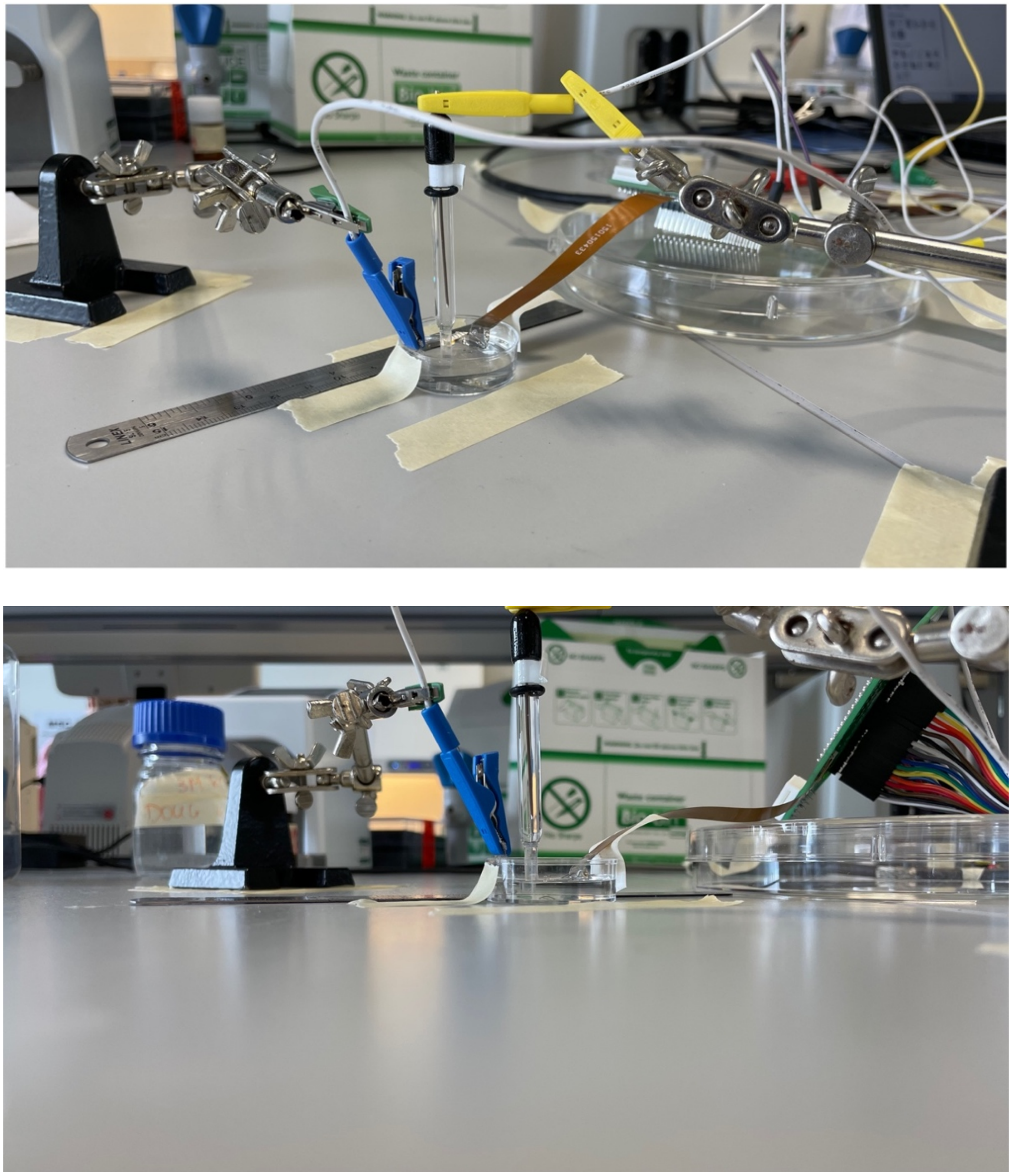


**Figure S3.** Image of electrical impedance spectroscopy (EIS) measurement setup (top). Blue clip goes to a platinum mesh acting as the counter electrode. Yellow clip goes to Ag/AgCl electrode acting as the reference. It is placed between the counter and the device. Device is on the right side of the setup, running out a to a custom PCB, designed to access the different channels on the device. All electrodes were submerged in PBS for the measurement (bottom).

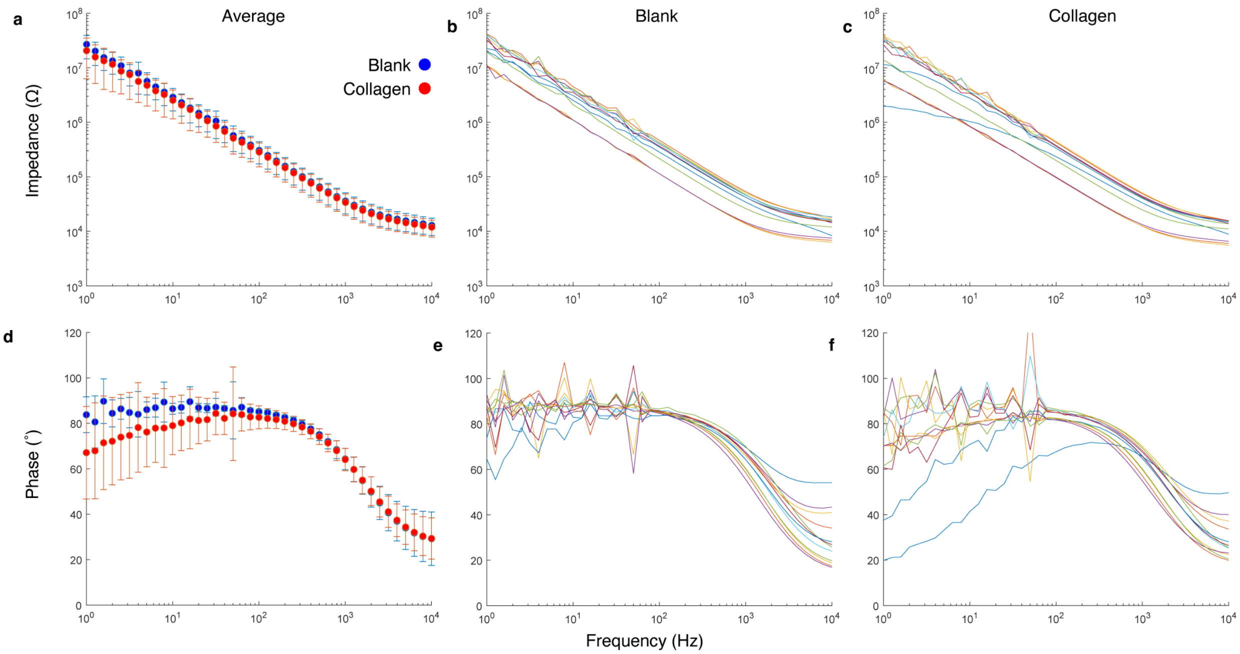


**Figure S4.** EIS measurements for implants, comparing impedance before and after collagen embedding for n = 12 arbitrary electrodes from N = 3 devices. Impedance spectra showing **a** average magnitude of impedance for all implants with and without collagen, **b** magnitude of impedance for individual electrodes without collagen and **c** individual electrodes after collagen embedding. Spectra showing **d** average phase of impedance for all implants with and without collagen, **e** phase of impedance for individual electrodes without collagen and **f** individual electrodes after collagen embedding. Colors in **b**, **c**, **e**, and **f** correspond to the same electrode for each plot. Plots **a** and **d** are the same as those in Figure 2e,f and are only shown here for comparison.

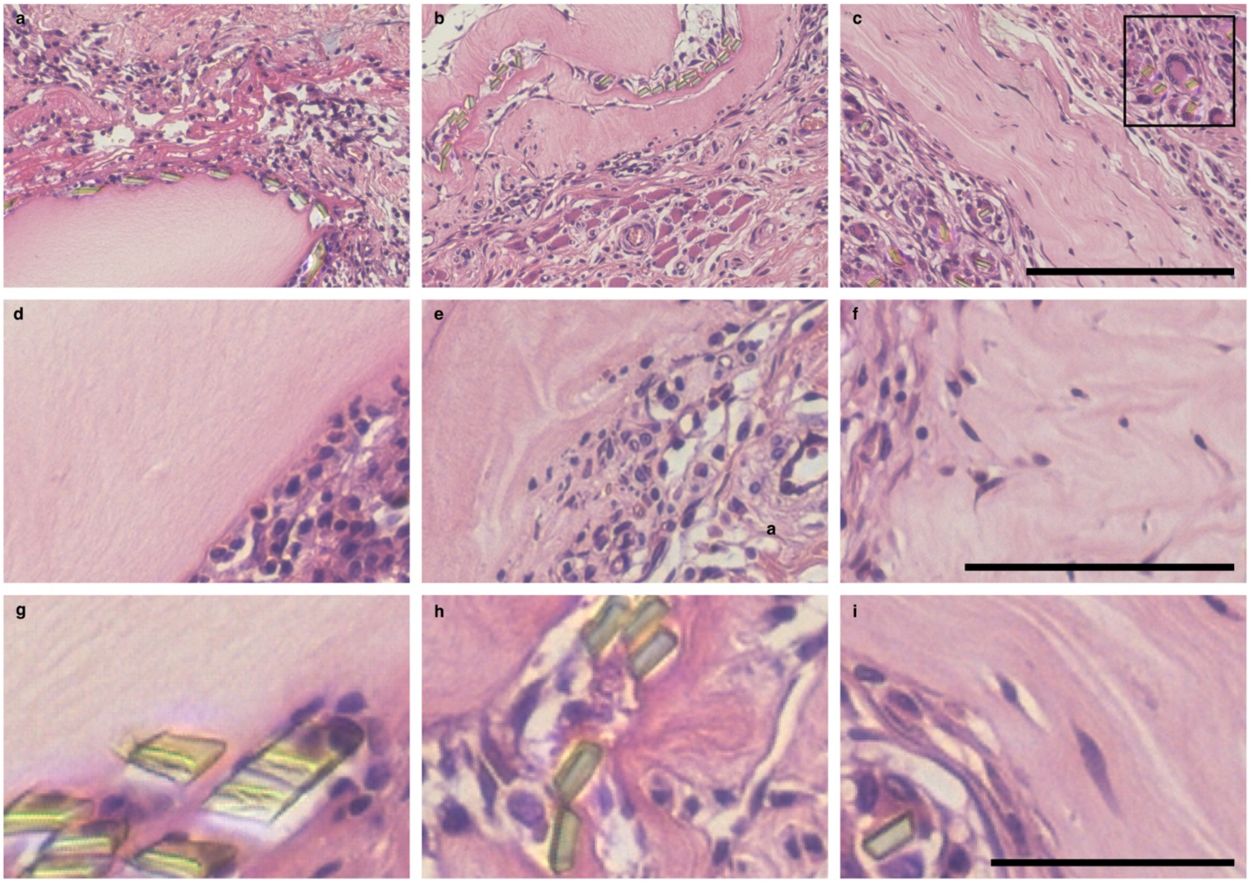


**Figure S5.** Hematoxylin and eosin images for hybrid implants at Days 3, 7, and 14 for examination of collagen behavior. Images of implant site at **a** Day 3, **b** Day 7, and **c** Day 14. Inset in **c** shows possible cellular encapsulation for one lead. Scale bar 200 mm and is the same for all three images. Images of collagen gels at **d** Day 3, **e** Day 7, and **f** Day 14. Scale bar is 100 mm and is the same for all three images. Images of implant lead cross-sections at **g** Day 3, **h** Day 7, and **i** Day 14. Scale bar is 50 mm and is the same for all three images. Dummy implants, lacking gold and PEDOT:PSS coatings were used for surgeries. Collagen gels were made at 10 mg/mL. Sections were prepared via paraffin embedding.

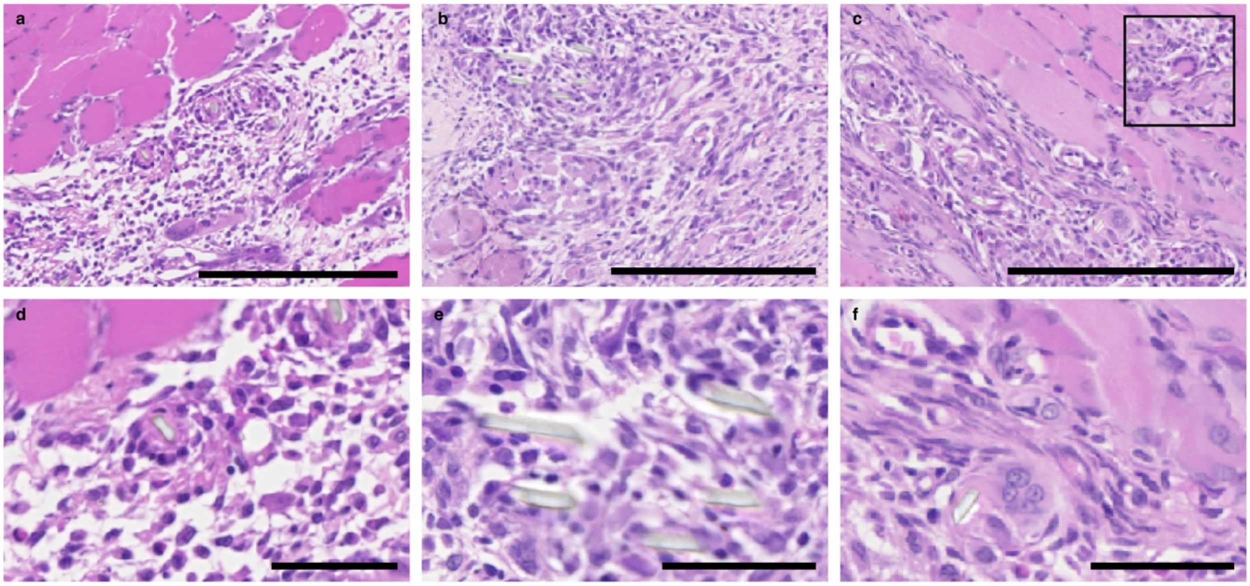

**Figure S6.** Hematoxylin and eosin images for implants lacking collagen at Days 3, 7, and 14. Images of implant site at **a** Day 3, **b** Day 7, and **c** Day 14. Inset in **c** shows possible cellular encapsulation for one implant lead. Scale bars are each 200 mm. Images of implant lead cross-sections at **d** Day 3, **e** Day 7, and **f** Day 14. Scale bars are each 50 mm. Dummy implants, lacking gold and PEDOT:PSS coatings were used for surgeries. Implants were placed used a 50 mm tungsten wire as an insertion shuttle. Sections were prepared via paraffin embedding.

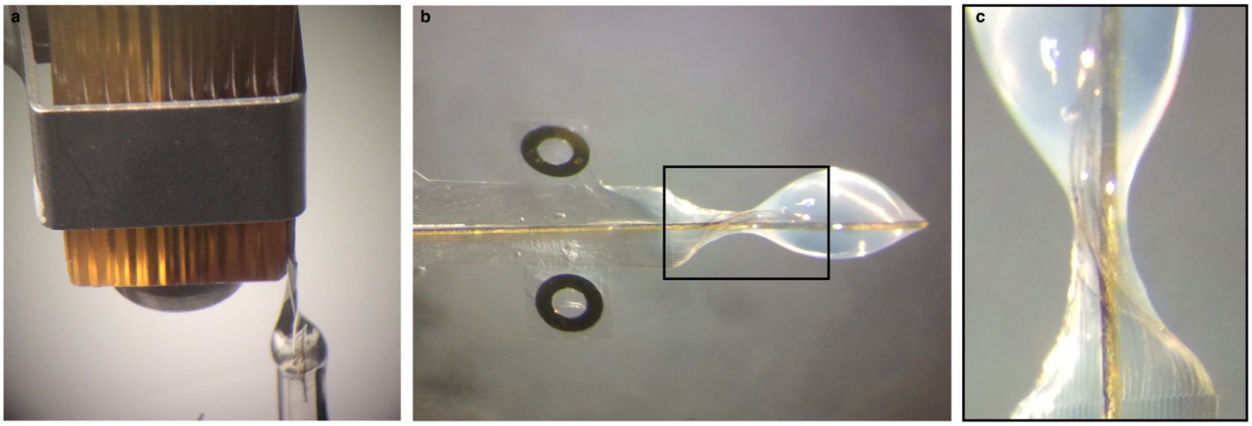

**Figure S7.** Image showing hybrid implant assembly. **a**, Image of implant gelling to create hybrid implant. **b**, Image of hybrid implant with embedded electrodes. **c**, Leads are visible distributed inside gel. This implant was constructed with a 50 mm diameter tungsten wire, which was used an insertion shuttle. This wire was removed after implantation for studies in which it was used.

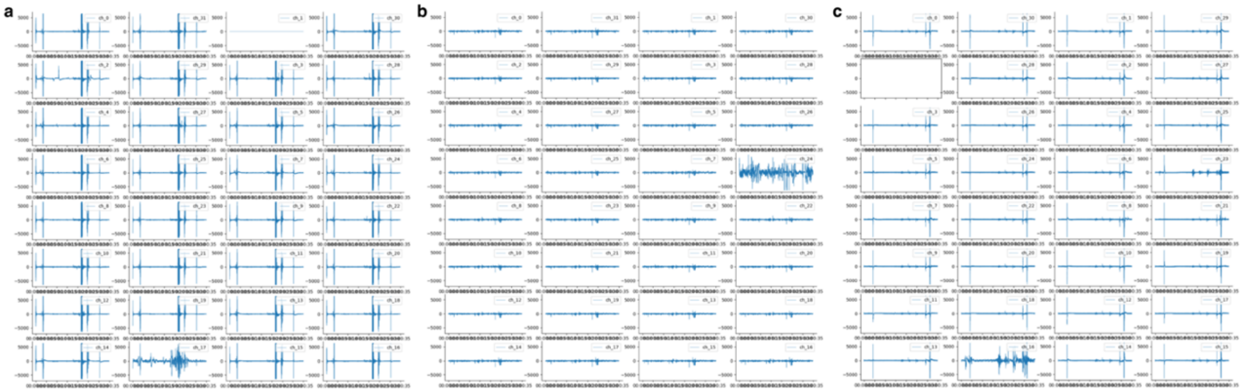

**Figure S8.** All channel recordings for **a** Day 1, **b** Day 3, and **c** Day 7.

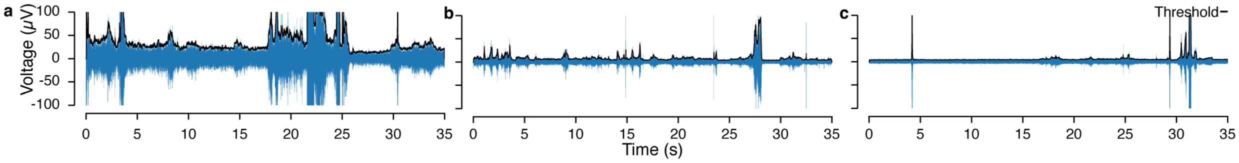


**Figure S9.** Threshold calculation example for **a** Day 1, **b** Day 3, and **c** Day 7 from Figure 4.

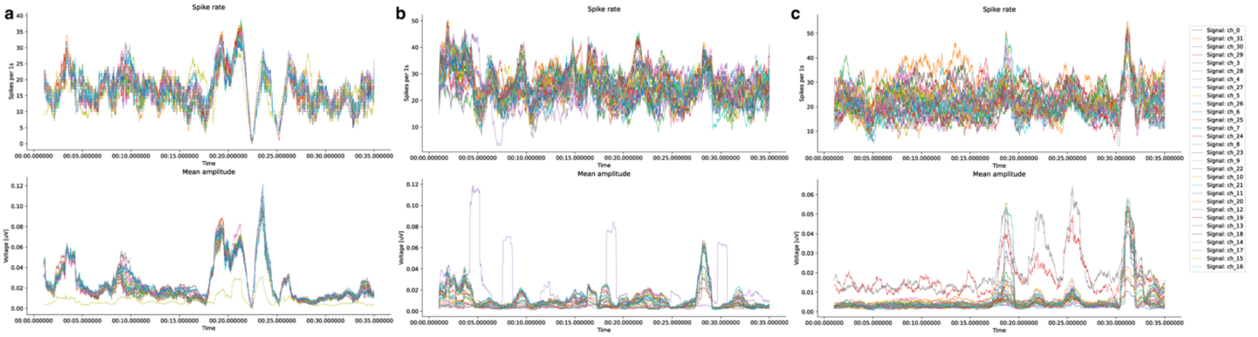


**Figure S10.** Spike rate (top) and mean spike amplitude (bottom) for all channels at **a** Day 1, **b** Day 3, and **c** Day 7.

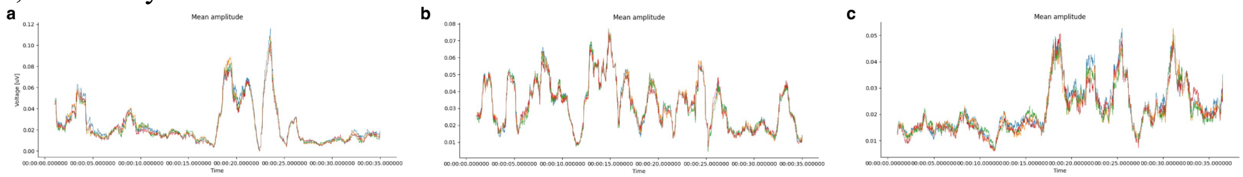


**Figure S11.** Example of tetrode set showing similar recordings for all tetrodes in set at **a** Day 1, **b** Day 3, and **c** Day 7.

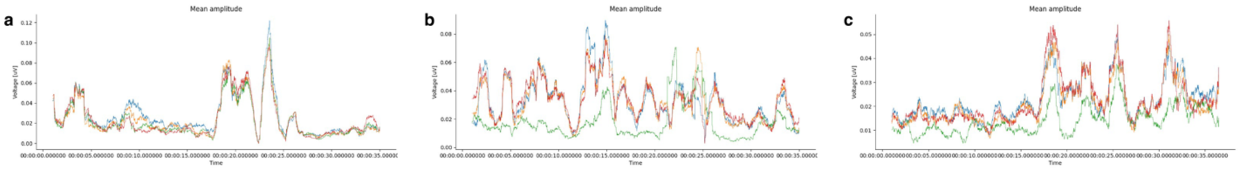

**Figure S12.** Examples of electrodes from other mixed tetrodes at **a** Day 1, **b** Day 3, and **c** Day 7.

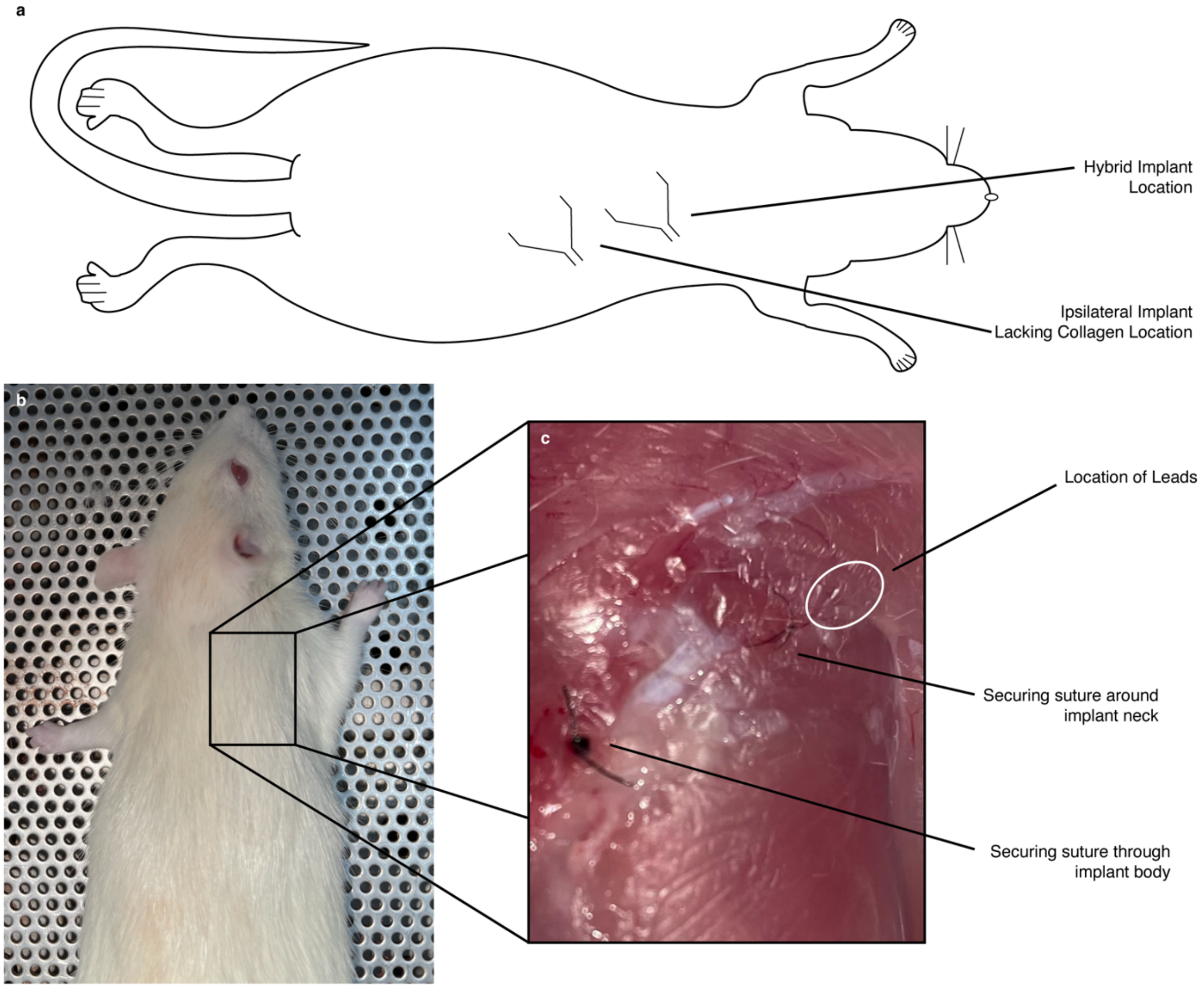


**Figure S13. a**, Schematic showing approximate location of implant with control. **b**, Image of gross appearance of tissue and implantation site for rat with hybrid implant after 2 months of implantation. **c**, Inset shows tissue at implantation site. Some fibrosis is visible near the suturing point at the back of the implant, as would be expected. Site containing implant leads is circled.

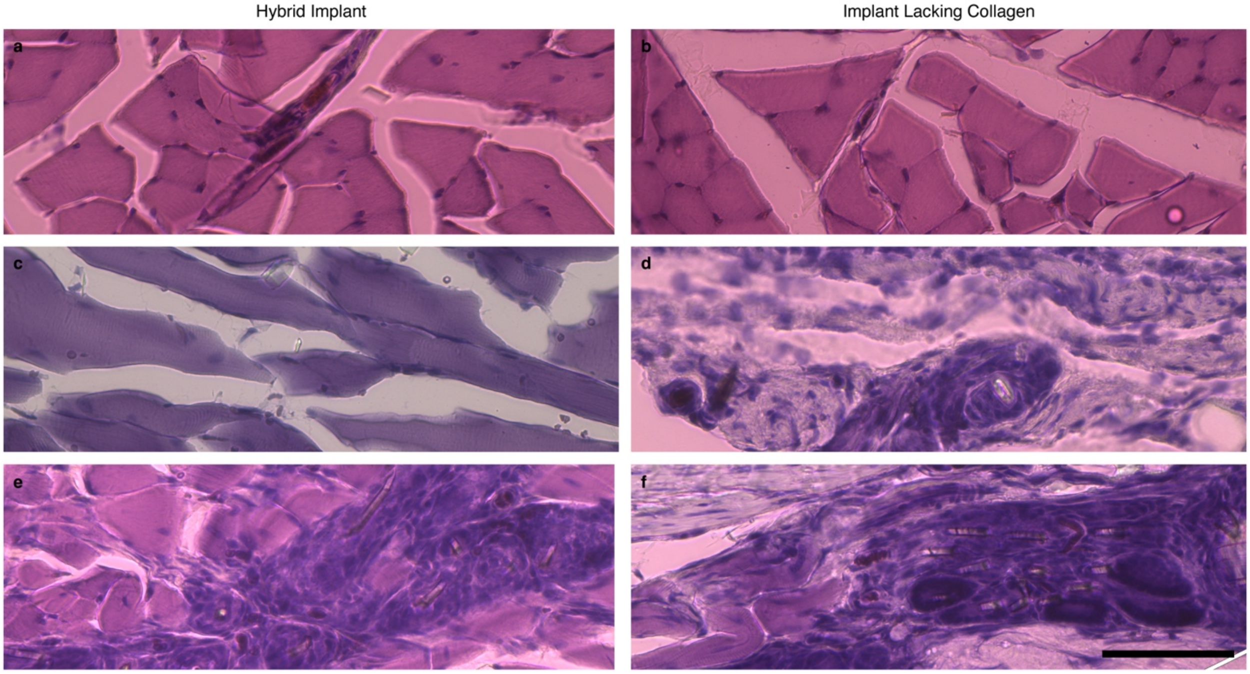


**Figure S14.** Histological images, stained with hematoxylin and eosin, showing leads, which were located for rat 1 (**a**, **b**), rat 2 (**c**, **d**), and rat 3 (**e**, **f**), with hybrid implants shown in the left column and implants lacking collagen shown in the right column. Scale bar is 100 mm and is the same for all images. We performed this surgery on 7 animals. We performed a dissection on one animal to examine the gross appearance of the tissue and determine preparation methods for histology. We were able to visualize the implanted leads in the tissue through this dissection, allowing us to designate methods for further sectioning. We sectioned tissue from the remaining animals (n = 6), but we were only able to locate leads in the sections for half of these animals. As the leads are approximately 10 x 4 mm in size, they are challenging to locate within the tissue during histological sectioning. However, we were able to identify the approximate location of the implants for all animals (n = 6) by examining the tissue for suture locations. By this means, we were able to look for remnant collagen. Sections were prepared via cryo-embedding.

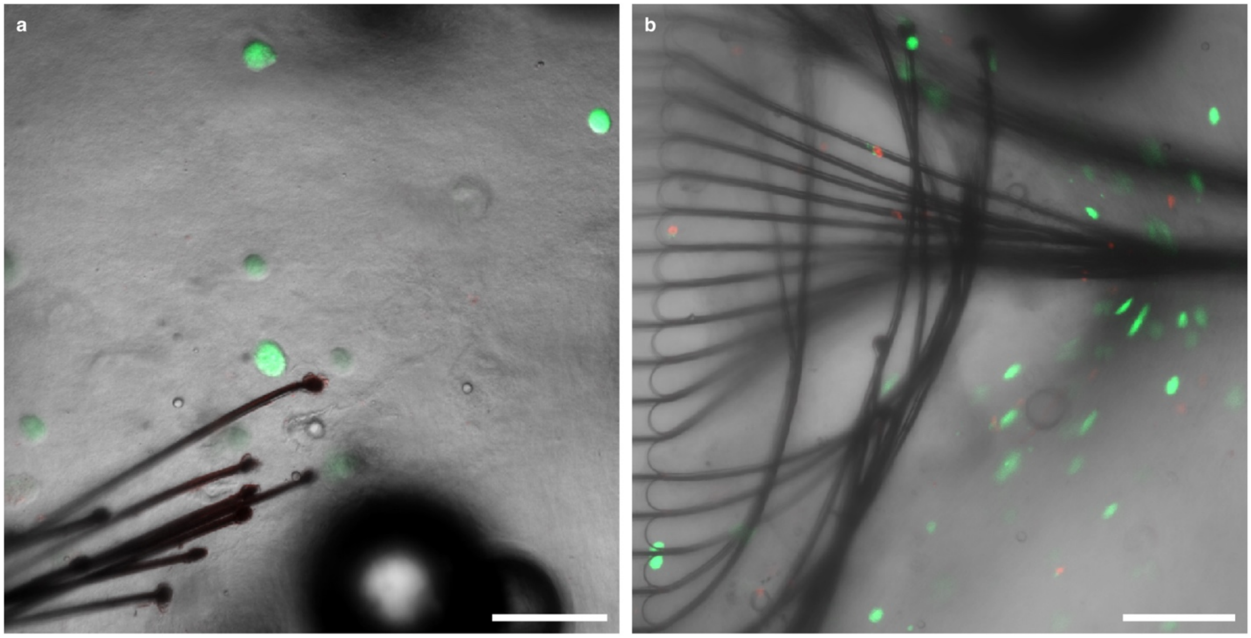


**Figure S15.** Images showing cellularized gel construction capabilities. **a**, Confocal projection from a z-stack (83 images across 82 mm) of rat bone marrow stromal cells in a collagen gel with embedded electrodes. We used bone marrow stromal cells, which is a common source for mesenchymal stem cells, as these cells have been shown to be useful for recapitulating various phenotypes in the musculoskeletal system. Cells were mixed into an 8 mg/mL gel at a final concentration of 250,000 cells/mL. **b**, Confocal image of SH-5YSY neuroblastoma cells in an 8 mg/mL gel with an embedded device at a final concentration of 500,000 cells/mL. Both images show cells stained with calcein AM and 1-ethidium homodimer immediately after injection to determine effect of hybrid implant fabrication on cell viability. Images were collected using Zeiss electronically switchable illumination and detection module (ESID). Scale bars for both images are 100 mm.

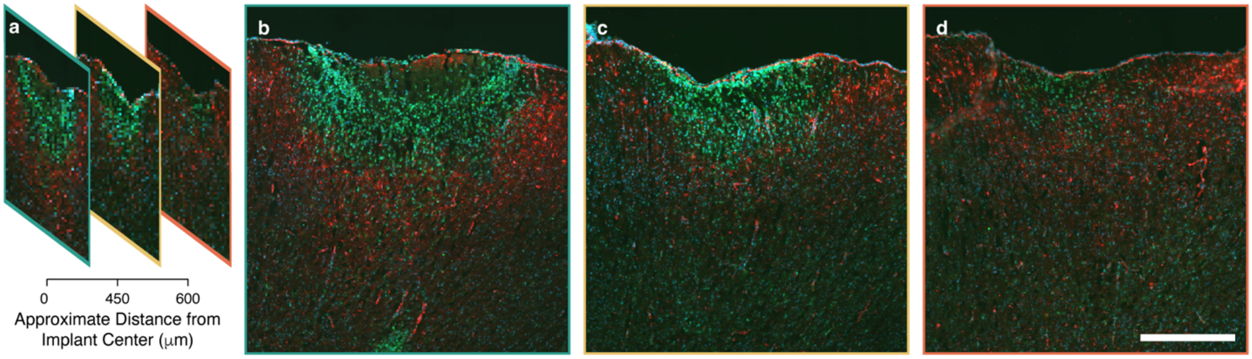


**Figure S16.** Cortical implantations of hybrid devices and localized immune response. **a**, Schematic of images showing approximate distance between sections moving from interior of implant to periphery. Immunofluorescence images of **b** implant interior, **c** implant edge, and **d** implant periphery. Samples were prepared via cryo-sectioning and stained for activated microglia (CD11b and CD11c positive – green), astrocytes (GFAP – red), and cell nuclei (blue). Scale bar is 500 mm. Dummy implants, lacking gold and PEDOT:PSS coatings were used for surgeries. Implants were placed with a 50 mm tungsten wire shuttle (Figure S7). Collagen gels were made at 8 mg/mL. Sections for brain analysis were prepared via cryo-embedding.

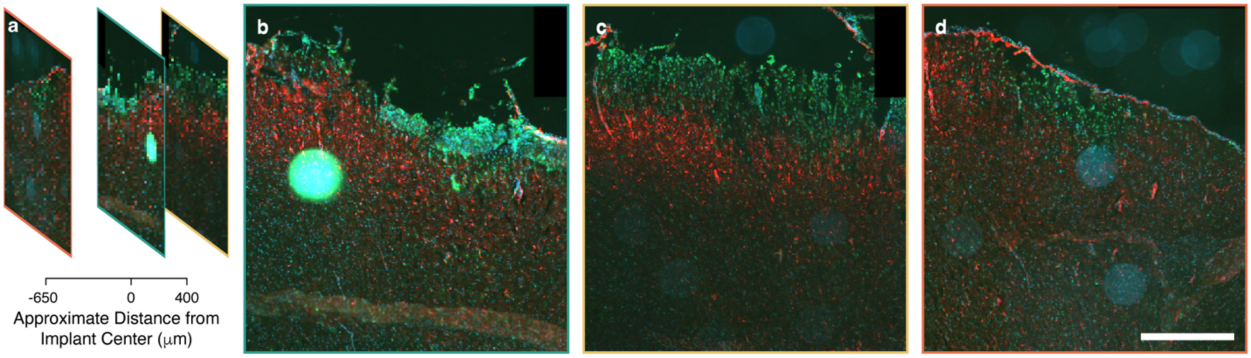


**Figure S17.** Immunofluorescence image of implant lacking collagen. **a**, Approximate location of each image. **b**, Image from approximate center of implant site. **c**, Image from ~400 mm away from center. **d**, Image from ~650 mm away from center. Note that images are arranged to match Figure S16 for comparison. Images **c** and **d** are at slightly different distances than in Figure S16 and are taken from opposite sides of the implant site, due to histological processing methods. Activated microglia (CD11b+CD11c positive) are green, astrocytes (GFAP) are red, and cell nuclei (DAPI) are blue. Scale bar is 500 mm. Dummy implants, lacking gold and PEDOT:PSS coatings were used for surgeries. Implants were placed with a 50 mm tungsten wire shuttle.

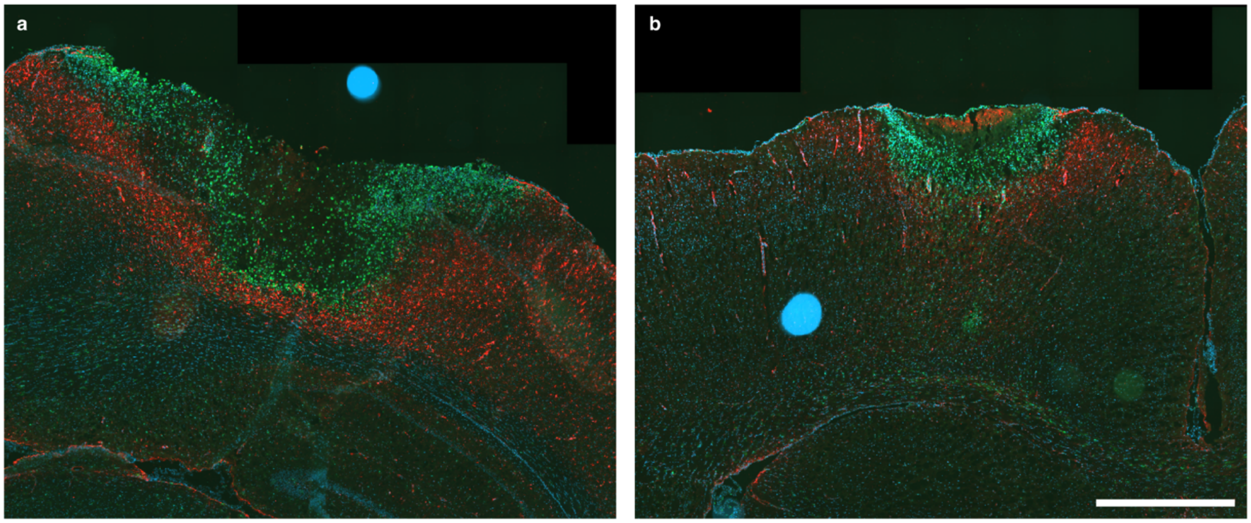


**Figure S18.** Immunofluorescence image of whole hemisphere for hybrid implants in n = 2 rats (**a**, **b**). Images show the approximate center of the implant for each rat. Activated microglia (CD11b+CD11c positive) are green, astrocytes (GFAP) are red, and cell nuclei (DAPI) are blue. Scale bar is 1 mm.

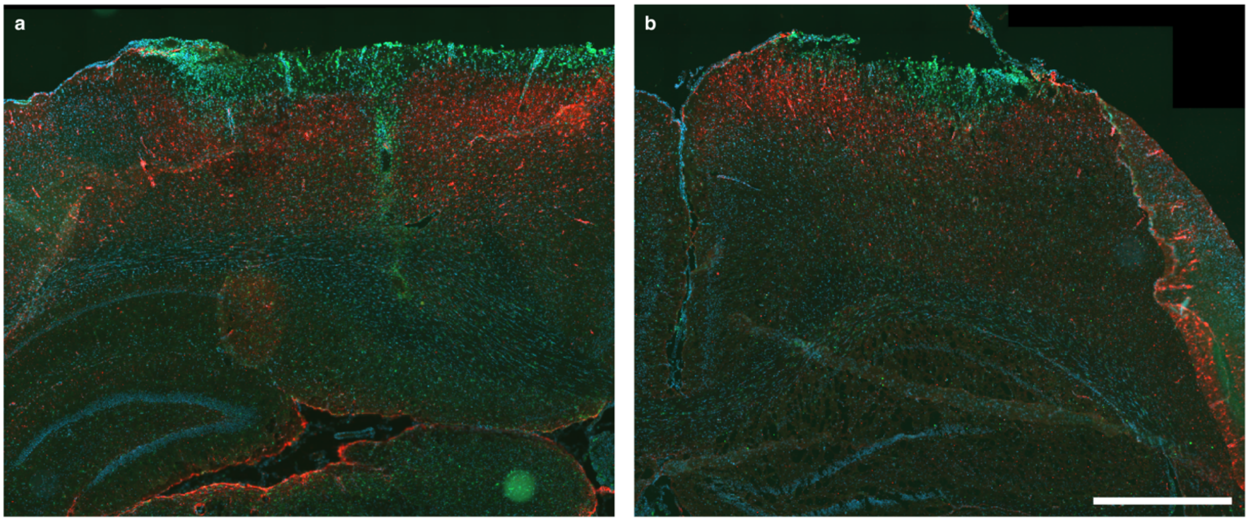


**Figure S19.** Immunofluorescence image of whole hemisphere for implants lacking collagen in n = 2 rats (**a**, **b**). Images show the approximate center of the implant for each rat. Activated microglia (CD11b+CD11c positive) are green, astrocytes (GFAP) are red, and cell nuclei (DAPI) are blue. Scale bar is 1 mm.

**Table S1.** Table summarizing histology results for 2-month intramuscular study. Rat 1 corresponds to Figure S14a,b, Rat 2 to Figure S14c,d, and Rat 3 to Figure S14e,f.

| | Foreign Body Response | |
|---|---|---|
| | Hybrid Implant | Implant Lacking Collagen |
| Rat 1 | | |
| Rat 2 | | x |
| Rat 3 | x | x |

**Video S1.** Video with paired time trace at Day 1. Data are bandpass filtered from 500 – 2000 Hz. The cage was fitted with a light that could be connected to a pulse generator on the recording device. By applying a pulse to the light, we could sync live videos with recordings.
